# SCON - A <u>S</u>hort <u>C</u>onditional intr<u>ON</u> for conditional knockout with one-step zygote injection

**DOI:** 10.1101/2021.05.09.443220

**Authors:** Szu-Hsien Sam Wu, Réka Szép-Bakonyi, Heetak Lee, Gabriele Colozza, Ayse Boese, Krista Rene Gert, Natalia Hallay, Jihoon Kim, Yi Zhu, Sandra Pilat-Carotta, Hans-Christian Theussl, Andrea Pauli, Bon-Kyoung Koo

## Abstract

The generation of conditional alleles using CRISPR technology is still challenging. Here, we introduce a Short Conditional intrON (SCON, 189 bp) that enables rapid generation of conditional alleles via one-step zygote injection. SCON has conditional intronic function in various vertebrate species and its target insertion is as simple as CRISPR/Cas9-mediated gene tagging.

## Introduction

CRISPR gene editing has facilitated the investigation of gene function by precise gene knockouts or knock-ins. Upon induction of gRNA-directed double-strand breaks (DSBs), the preferred repair pathway, non-homologous end-joining (NHEJ), often leads to random insertions or deletions (indels), where out-of-frame mutations can cause partial or complete loss of gene function. On the other hand, by using a DNA template, DSBs can also be repaired via homology-directed repair (HDR), which allows precise knock-ins of various sequences into target loci. Due to the versatility and wide applicability of the CRISPR/Cas9 system, it has been utilized in numerous cell lines and lab organisms, spanning the entire biological and biomedical research fields^1^.

Despite the revolutionary advancement of CRISPR technology, the generation of conditional alleles has not been as easy as that of knockout or knock-in alleles. A conditional knockout (cKO) approach is often required to study essential genes such as housekeeping or developmentally required genes, as it allows spatiotemporal control of the gene knockout, thereby avoiding the early lethality associated with a simple knockout. The use of cKO for rodents, and eventually for non-human primates, therefore, also contributes to improved animal welfare. For many years, the Cre/LoxP system has been widely utilized to construct cKO alleles, by inserting two LoxP recombination sites into the introns flanking essential exon(s). The generation of such “floxed” alleles in mice has traditionally involved the use of mouse embryonic stem (ES) cells, which are microinjected into blastocysts to generate chimeric mouse embryos^2^. Recently, cKO alleles have also been generated via CRISPR/Cas9-mediated insertion of LoxP sites in zygotes. However, this turned out to be rather challenging, even with additional refinements^3^.

Here, we introduce the use of a universal conditional intron system for cKO approaches suitable for various animal models. Such conditional intron approach has been attempted in the past, as it enables simple insertional mutagenesis with a fixed universal conditional intronic cassette^4–6^, but it has not been widely utilized in animal models because the cassettes were either too long^4,5^ or led to unexpected hypomorphic effects^6^.Notably, presently the simplest form is DECAI^6^. Although its short size (201 bp) makes it desirable for gene targeting via zygote injections, it reduces target gene expression that compromises normal gene expression and animal development (data not shown). Here, we present a fully optimized Short Conditional intrON (SCON) cassette that shows no hypomorphic effects in various vertebrate species and is suitable for targeting via one-step zygote microinjection.

### SCON is a versatile conditional intron with no discernable hypomorphic effect

SCON is a modified intron derived from the first intron (130 bp) of the human *HBB* (hemoglobin subunit beta) gene. SCON is 189 bp long, consisting of, from the 5’ to 3’ end, a splice donor, a LoxP recombination site, a branch point, a second LoxP recombination site, a polypyrimidine tract, and a splice acceptor. By design, it has a similar sequence architecture to the previously introduced conditional intronic system DECAI (201 bp)^6^, which showed a hypomorphic effect at the level of protein expression. To optimize conditional intron function, several features were implemented into SCON: 1) the length between the putative branch point and the splice acceptor was kept to 45 bp to allow efficient splicing to take place, 2) 100 bp distance was used between the two LoxP sites for efficient Cre-LoxP recombination, and 3) miscellaneous changes were incorporated for optimal splice donor, acceptor and pyrimidine tract sequences. Upon recombination, SCON is reduced to 55 bp, of which all three reading frames contain translational stop codons within the remaining LoxP sequence, such that loss of gene function occurs via premature translational termination (Fig. 1a).

**Fig.1.**
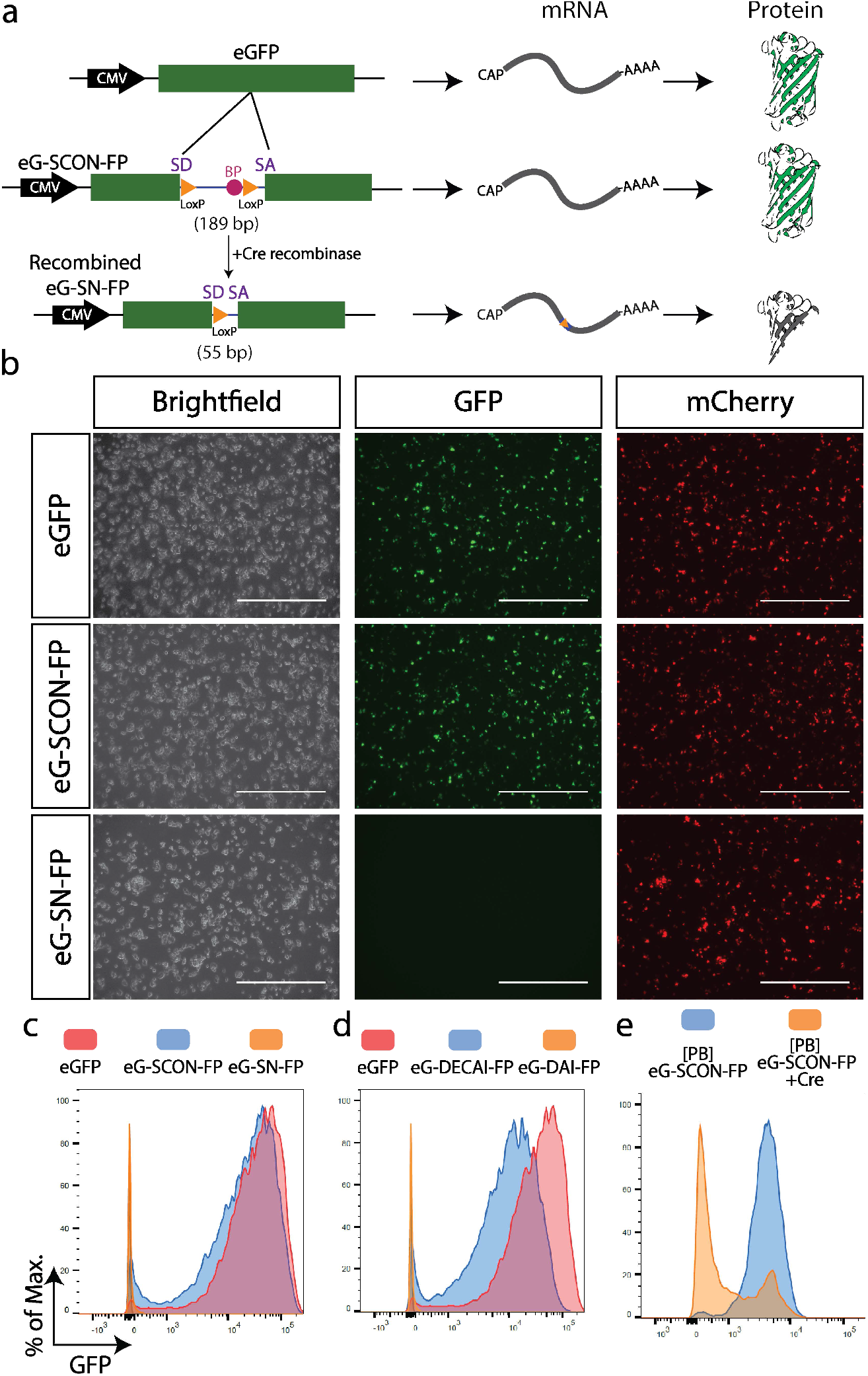
SCON functionality test with an eGFP overexpression construct. **a**, Schematic diagram of SCON functionality test in an eGFP overexpression construct including intact eGFP, eG-SCON-FP and recombined eG-SN-FP. SD, splice donor; BP, branch point; SA, splice acceptor. **b**, Images of transfected HEK293T cells on day 1, with intact eGFP, eG-SCON-FP and recombined eG-SN-FP. Scale bar, 1 mm. All constructs were co-transfected with an mCherry-overexpression plasmid. **c, d**, Histograms of the flow cytometry analysis of transfected HEK293T cells showing comparisons between eGFP (red) and eG-SCON-FP (blue) (**c**), between eGFP (red) and eG-DECAI-FP (blue) (**d**), and the respective recombined forms eG-SC-FP and eG-DAI-FP (yellow) (**c**,**d**). **e**, Flow cytometry analysis of mouse ES cells with integrated piggyBac-eG-SCON-FP transfected with Cre-expressing plasmid (yellow) or empty vector (blue).

Firstly, we validated the functionality of SCON in an eGFP overexpression construct. We co-transfected HEK293T cells with an mCherry cassette (serving as transfection control) and a cassette containing either the intact eGFP, eG-SCON-FP or already Cre-recombined eG-SCON-FP (eG-SN-FP) (Fig. 1a), and assessed the fluorescence intensity by fluorescent microscope and flow cytometry (Fig. 1b, c). We also carried out the same test in parallel with DECAI for comparison. Both eG-SCON-FP and eG-DECAI-FP showed GFP expression, whereas the recombined form had no detectable fluorescence (Fig. 1b-d). Interestingly, eG-SCON-FP exhibited similar levels of GFP signal as the intact eGFP construct (Fig. 1c). However, eG-DECAI-FP showed reduced levels (Fig. 1d), indicating an adverse hypomorphic effect as an intron, which is in-line with the previous observations^6^. Thus, our SCON showed a reliable conditional intronic function with no discernable hypomorphic effect.

To better understand the functionality of different parts within the SCON cassette, we designed a series of “A-stretch variants” in which 6-10 nt within SCON were converted to adenine (Extended Data Fig. 1a). Out of the 13 different variants, we found that four showed either hypomorphic or complete reduction of eGFP fluorescence when inserted as an intron (Extended Data Fig. 1b, c). These included the branch point (variant 11) and the polypyrimidine tract (variant 12), which resulted in hypomorphic expression of the inserted eGFP cassettes, whereas the splice donor (variant 1) and the splice acceptor (variant 13) resulted in complete loss of eGFP expression (Extended Data Fig. 1b, c). These data suggest that all elements for splicing are required for optimal intronic function of SCON, while most parts between the two LoxP sites are less essential.

Next, we tested whether SCON can be efficiently recombined in mammalian cells. We cloned the eG-SCON-FP construct into a piggyBac transposon backbone, and generated a clonal mouse ES cell line with constitutive eGFP expression containing SCON. Co-transfecting a Cre-expressing plasmid with an mCherry cassette into these eG-SCON-FP-expressing ES cells resulted in efficient reduction of eGFP levels within the mCherry+ population, whereas the mock control (transfected only with the mCherry cassette) maintained high levels of eGFP (Fig. 1e). Taken together, our data indicate the suitability of SCON for use in mammalian systems, where it is neutral upon insertion and can be efficiently recombined by Cre recombinase to abolish its intronic function and cause knockout of the inserted gene.

### Neutrality of SCON is conserved in various vertebrate species

cKO alleles have been widely used in mice for decades thanks to the robust methodology using germline-competent ES cells^7^, the endeavor of the International Knockout Mouse Consortium^8^, and, recently, the advent of CRISPR technology^9^. However, for many other vertebrate species such as zebrafish, frog, rat, porcine, bovine and non-human primate models, the use of the cKO approach has been limited, mainly due to the lack of reliable germline-competent ES cells. We therefore sought to test whether SCON may be a suitable cKO strategy for non-murine species. To this end, we used cell lines from different species, including C6 (*Rattus norvegicus*), PK15 (*Sus scrofa*), LLC-MK2 (*Macca mulatta*) and Vero (*Cercopithecus aethiops*) and transfected them with overexpression constructs of eGFP, eG-SCON-FP or eG-DECAI-FP and the corresponding Cre-recombined forms. In line with the results in HEK293T and mouse ES cells, the SCON intron did not cause any discernable hypomorphic effects in any of the cell lines tested (Extended Data Fig. 2a-d, in blue), while the recombined forms of SCON abrogated the eGFP expression completely (Extended Data Fig.2 b-e, in orange). Intriguingly, in three of the cell lines (C6, PK15 and Vero) eG-DECAI-FP showed reduced levels of eGFP compared to eG-SCON-FP (Extended Data Fig. 2e), again confirming that SCON is devoid of the hypomorphic effects associated with DECAI. In addition, we also explored the possibility of utilizing SCON in frog (*Xenopus laevis*) and zebrafish (*Danio rerio*) by injecting the eGFP, eG-SCON-FP and the Cre-recombined form plasmids into 4-cell stage embryos or fertilized eggs, respectively. We examined the embryos 24 hours post-injection and observed that eGFP and eG-SCON-FP expressed readily detectable eGFP fluorescence whereas the embryos that received the recombined form did not (Extended Data Fig. 2f, g). Together, these data indicate that SCON has the potential to be utilized as a cKO system in a wide variety of vertebrate species.

### Targeted insertion of SCON via one-step zygote injection

As our results from the *in vitro* overexpression-based systems demonstrated the suitability of SCON for cKO approaches, we next sought to test whether it would also work well for targeting endogenous genes *in vivo*. Therefore, we chose to generate a SCON cKO *Ctnnb1* allele (Extended Data Fig. 3a), which encodes for β-catenin and is a developmentally required gene with early lethality upon knockout in mice. We injected CRISPR ribonucleoprotein (RNP) with 300 bp-long, commercially-synthesized, single-stranded deoxynucleotides (ssODNs) consisting of SCON with short homology arms (55 and 56 bp for the 5’ and 3’ homology arms, respectively) into the cytoplasm of developing 2-cell stage mouse embryos^10^. Genotyping and sequencing revealed that one out of the resulting 13 offspring (7.7%) had a precise heterozygous integration with the other allele remaining intact (Extended Data Fig. 3b). From this offspring, we were able to backcross to confirm germline transmission, and breed to homozygosity (*Ctnnb1*^sc/sc^) (Extended Data Fig. 3c). From the heterozygote to heterozygote (HET) crosses, we did not observe under-represented ratios of homozygous (HOM) mice (Extended Data Fig. 3d) and the HOM mice showed no discernible phenotype, confirming intact gene function with SCON insertion. Similarly, we generated a second SCON cKO *Sox2* allele with frt recombination sites (Extended Data Fig. 3e). Sox2 mice appear normal and could be bred to homozygosity (Extended Data Fig. 3f) and HET-HET crosses yielded expected proportion of HOM mice (Extended Data Fig. 3g), again indicating that SCON is suitable for generating cKO alleles *in vivo*.

To verify the conditional functionality of the *Ctnnb1*^*sc*^ allele, we utilized the *Villin-CreER* ^*T2*^ (*Vil-CreER* ^*T2*^) for intestinal epithelium-specific Cre recombination. We first isolated crypts from the duodenum of HOM (*Vil-CreER*^*T2*^; *Ctnnb1*^sc/sc^) and wild-type (*Ctnnb1*^+/+^) mice to establish adult stem cell-based intestinal organoids. Then, budding organoid cultures (with EGF, Noggin and R-spondin) were transiently treated with 4-hydroxytamoxifen (4-OHT) for 8 hours. Both 4-OHT treated and untreated wild-type organoids as well as untreated HOM organoids continued to grow normally (Fig. 2a). However, 4-OHT treated HOM organoids ceased growth or collapsed from day 2 onward. Immunolabelling confirmed the loss of β-catenin in the small cystic mutant organoids, while DAPI and phalloidin staining shows that the cells remain live (Fig. 2a).

**Fig 2.**
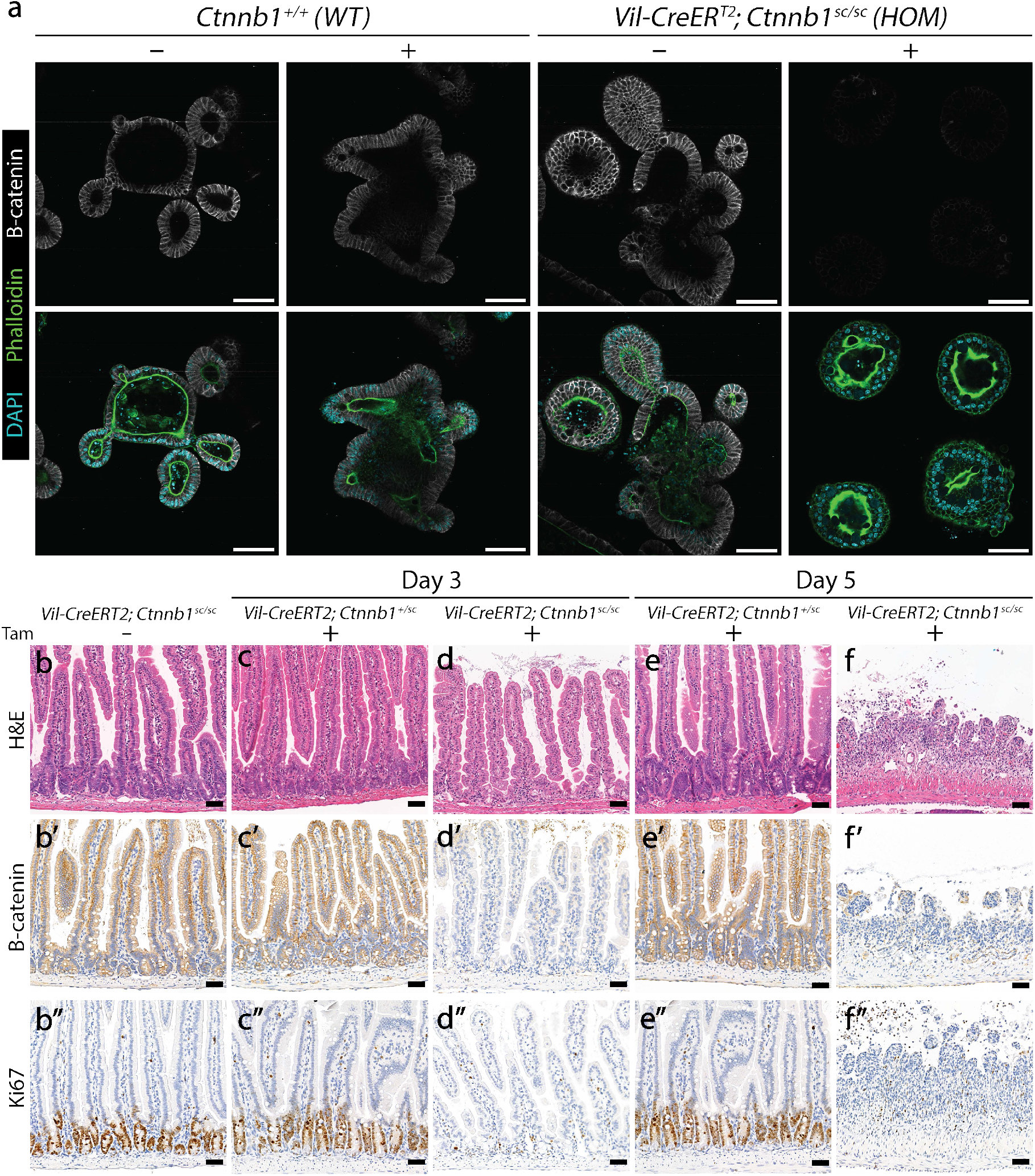
*Ctnnb1*^*sc*^ mice harbor a functional conditional allele that works *in vivo*. **a**, Small intestinal organoids from WT (*Ctnnb1*^*+/+*^) and HOM (*Vil-CreER*^*T2*^; *Ctnnb1*^*sc/sc*^), treated with either 4-OH-tamoxifen (4-OHT) or vehicle for 8 hours. Organoids were fixed on day 4 and stained for β-catenin (grey), phalloidin (green) and DAPI (cyan). Scale bar, 100 μm. Homozygous (*Vil-CreER*^*T2*^; *Ctnnb1*^*sc/sc*^) intestines are healthy, with normal epithelial crypt-villus morphology (**b**), β-catenin on the cell membrane (**b’**) and Ki67 marking proliferating cells in the crypts (**b’’**). Heterozygous (*Vil-CreER*^*T2*^; *Ctnnb1*^*+/sc*^) intestines are unaffected in morphology (**c, e**), β-catenin (**c’, e’**) and Ki67 (**c’’, e’’**) after tamoxifen treatment. In homozygous intestines, tamoxifen treatment leads to loss of crypts (**d**), β-catenin (**d’**) and Ki67 (**d’’**) staining on day 3. On day 5, the epithelium was completely lost (**f**-**f’’**). H&E, hematoxylin and eosin. Scale bar, 50 μm.

To directly verify the functionality *in vivo*, we injected 3 mg tamoxifen (TAM) per 20 g body weight into both HOM (*Vil-CreER* ^*T2*^; *Ctnnb1*^sc/sc^) and HET (*Vil-CreER* ^*T2*^; *Ctnnb1*^+/sc^) mice, and harvested the intestines on day 3 and day 5. Control samples showed a normal crypt-villus axis, detectable membrane-bound β-catenin, and a Ki67+ proliferative zone in the bottom of the crypts where stem and progenitor cells are located (Fig. 2b-b’’, c-c’’, e-e’’). On day 3, TAM-treated HOM samples showed clear loss of proliferative crypts with the loss of β -catenin staining (Fig. 2d-d’’). On day 5, TAM-treated HOM mice had shortened and inflamed intestines, sections of which showed nearly complete loss of intestinal epithelium (Fig. 2f-f’’). These data indicate efficient recombination of the *Ctnnb1*^sc^ allele *in vivo* and the loss of β-catenin function upon recombination.

### SCON is applicable in large fraction of protein-coding genes

To systematically estimate targetable sites for SCON, we carried out bioinformatic analysis to screen for possible insertion sites in mouse, rat, macaque, marmoset and medaka genomes (Extended Data Fig. 5). The selection criteria include: 1) target exons are positioned within the first 50% of the protein coding-sequence from canonical transcripts of protein-coding genes; 2) intron insertion sites contain either stringent (MAGR, (A/C)-A-G-(A/G))^11^ or flexible (VDGN, (A/G/C)-(A/T/G)-G-(A/T/G/C))^12^ splice junction consensus sequences; 3) exon must be larger than 120bp in length and that both 5’ and 3’ split exons to be at least 60bp (Extended Data Fig. 4a)^5^. After combining all selection criteria, we identified that majority of coding genes are targetable in all five species, on average 80.8% for MAGR and 87.7% for VDGN intron insertion sites (Extended Data Fig. 4b, c). We also found CRISPR/Cas9 targeting stie(s) around intron insertion sites in most cases (Extended Data Fig. 4c; Supplementary tables 1-10). This is a conservative estimate as some genes with an important domain close to the 3’ end can still be targeted in the second half and, if necessary, a novel intron insertion site can be generated by introducing silent mutations.

## Discussion

In summary, our SCON-mediated cKO approach transforms the complicated process of generating conditional alleles into a simple, CRISPR-mediated knock-in of a 189 bp intronic sequence through one-step zygote injection. This novel strategy therefore opens up the exciting possibility of generating conditional knockouts in other vertebrate models ranging from fish to non-human primates. A key to our success is the creation of a neutral artificial intron ready for Cre-mediated inactivation. In contrast to DECAI, which shows undesirable hypomorphic effects that compromise its utility in cKO animal models; SCON is well-tolerated and ideal for rapid generation of cKO animals. Moreover, the LoxP sequences can also be replaced by other recombination sites (e.g. FRT) for rapid generation of FRT- or other recombinase-based conditional alleles that are not yet widely utilized. Lastly, the dispensable region between the two LoxP sites serves as a harboring space for the addition of other genetic elements. We expect that the SCON strategy will become the foundation for a new cKO approach in biomedical and industrial research that is well suited for meeting animal welfare concerns.

## Materials and Methods

### Mice

All animal experiments were performed according to the guidelines of the Austrian Animal Experiments Act, and with valid project licenses approved by the Austrian Federal Ministry of Education, Science and Research and monitored by the institutional IMBA Ethics and Biosafety department.

#### Generation of Ctnnb1-SCON and Sox2-SCON mice

The new *Ctnnb1-SCON* (*Ctnnb1*^*sc*^) and *Sox2-SCON* (*Sox2*^*sc*^) conditional KO mouse lines were generated via 2-cell and zygote stage embryo injection. We prepared 25 μl of CRISPR injection mix in nuclease-free buffer (10 mM TRIS-HCl, pH 7.4 and 0.25 mM EDTA) consisting of spCas9 mRNA (100 ng/μl), spCas9 protein (50 ng/μl), sgRNA (50 ng/μl) and ssODN (20ng /μl, Genscript). The mixture was spun down in a tabletop centrifuge at 13,000g at 4 °C for 15-20 minutes to prevent clogging of the injection needles. Frozen 2-cell stage embryos of C57Bl/6JRj background (JANVIER LABS) were used for the cytoplasmic injection.

#### Tamoxifen administration and organ harvest

Ctnnb1-SCON was crossed with Vil-CreER^*T2*^ (JAX, 020282)^13^ and bred to obtain either HET (Vil-CreER^*T2*^; Ctnnb1^+/sc^) or HOM (Vil-CreER^*T2*^; Ctnnb1^sc/sc^) mice. Tamoxifen (Sigma, T5648) dissolved in corn oil (Sigma, C8267) or corn oil only was injected intraperitoneally into 8-12 week-old mice at a final concentration of 3 mg tamoxifen per 20 g body weight. Experiments were carried out in 2 mice of the respective genotype for each time point. Control mice received a matched volume of corn oil. On day 3 or day 5 mice were euthanized by cervical dislocation, and the intestines were harvested. The intestines were immediately cleaned with 1x PBS, flushed gently with 10% formalin solution (Sigma, HT501128), and fixed as ‘swiss rolls’ for 24 hours at room temperature. Fixed intestines were washed three times with 1x PBS, with 2-3 hours between each wash, before further processing.

### Genotyping

The toe clips or ear notches from the offspring were lysed in 30 μl of DirectPCR Lysis Reagent (Viagen) with 1 μl of proteinase-K (20 mg/ml; Promega, MC5005) at 55 °C overnight. The resulting mixture was diluted with 270 μl of nuclease-free water and spun down for at least 5 minutes in a tabletop centrifuge at 13,000 g. Then, 2-3 μl of the clear part of the solution was used for PCR with either Gotaq (Promega, M7808) or LongAmp 2X (NEB, M0287S), according to the manufacturer’s instruction. To check the sequences of genomic DNA, PCR bands of expected sizes were purified with a purification column.

### eGFP-SCON/ eGFP-DECAI constructs

The eG-SCON-FP and eG-DECAI-FP cassettes were synthesized and ordered from Genscript, and subsequently cloned into the pcDNA4TO construct with BamHI (R0136S, NEB) and XhoI (R0146S, NEB) via ligation with T4 ligase (M0202S, NEB). The vectors were recombined with Cre-expressing bacteria (A111, Gene bridges) to obtain the recombined forms. The correct clones were confirmed with restriction digest SalI (R0138S, NEB) and Sanger sequencing.

### SCON A-stretch variants inserted in the eGFP cDNA

eGFP cDNA containing the SapI recognition sites at the selected intron insertion site was synthesized and ordered from Genscript and cloned into pcDNA4TO construct with BamHI and XhoI. Different SCON variant fragments containing the respective complementary ends were then inserted into eGFP with SapI (R0569S, NEB) and T4 ligase for 20 cycles of 2 minutes at 37 °C and 5 minutes at 16 °C, followed by 15 minutes at 37 °C and 10 minutes at 80 °C incubations^5^. The mixture was transformed into *Escherichia coli* and DNA was extracted from individual colonies and checked with restriction digests and Sanger sequencing.

### Cell culture and transfection

#### HEK 293T cells

Human embryonic kidney (Hek) 293T cells were cultured in high glucose DMEM containing 10% fetal bovine serum (FBS, Sigma), 1% penicillin-streptomycin (P/S; Sigma, P0781) and 1% L-glutamine (L-glut; Gibco, 25030024).

#### Mouse ES cells

The mouse ES cell line AN3-12 was cultured as previously described^14^, in high glucose DMEM (Sigma, D1152) containing 10% FBS (Sigma), 1% P/S, 1% L-glut, 1% NEAA (Sigma, M7145), 1% sodium pyruvate (Sigma, S8636), 0.1 mM 2-mercaptoethanol (Sigma, M7522), 30 μg/ ml of mouse LIF (stock concentration: 2 mg/ ml).

#### Cell lines of other species

The following cell lines were cultured with basal medium supplemented with 10% FBS and 1% P/S. The basal medium for each cell line is indicated in brackets: C6 (ATCC, CCL-107; DMEM-F12 (Gibco, 31330038)), PK15 (Elabscience Biotechnology, EP-CL-0187; Minimal essential medium (Gibco, 11095080)), LLC-MK2 (Elabscience Biotechnology, ELSEP-CL-0141-1; RPMI-1640 (Sigma, R8758)), Vero (ATCC, CCL-81; DMEM-High glucose (Sigma, D1152)).

#### Plasmid transfection

500,000-750,000 cells were seeded in 6-well plates and left to attach and grow overnight. 2.5 μg of DNA (1μg of mCherry-expressing plasmid (Addgene, 72264), and 1.5 μg of pcDNA4TO-eGFP, -eG-SCON-FP, -eG-DECAI-FP or recombined forms of eG-SCON-FP or eG-DECAI-FP) was mixed with 8 μl of polyethyleneimine (1 mg/ ml; Polysciences, 23966) and incubated at room temperature for at least 15 minutes before being added dropwise to the cells. Culture medium was exchanged 8-10 hours after transfection. 36 hours after transfection, cells were examined under an EVOS M7000 microscope (Thermo Scientific) with the brightfield, GFP and TexasRed filters. 36-48 hours after transfection, cells were dissociated into single cells for flow cytometry analysis, with a BD-LSRFortessa flow cytometer (BD). Data from the flow cytometry experiments were analyzed in FlowJo software (BD). The fluorescence intensity values were exported, which were used for plotting and statistical tests using R.

### *X. laevis* and *D. rerio* embryo injection

*Xenopus laevis* eggs were collected and fertilized *in vitro*, as previously described^15^. 100 pg of DNA in total, consisting of 50 pg of pcDNA4TO-eGFP, pcDNA4TO-eG-SCON-FP or pcDNA4TO-eG-SN-FP, and 50 pg of mCherry-expressing plasmid, were injected into two dorso-animal blastomeres of *X. laevis* embryos at the 4-cell stage and imaged roughly 24 hours later, at early tailbud stage. For zebrafish experiments, 12 pg of DNA in total, consisting of 6 pg of pcDNA4TO-eGFP, pcDNA4TO-eG-SCON-FP or pcDNA4TO-eG-SN-FP, and 6 pg of mCherry-expressing plasmid, were injected into each fertilized egg/ 1-cell stage *D. rerio* embryo and imaged roughly 24 hours after. *X. laevis* embryos were imaged using a stereomicroscope (Leica, M165MC) equipped with GFP (excitation: ET470/40 nm; emission: ET525/50 nm) and mCherry (excitation: ET560/40 nm; emission: ET630/75 nm) filters. *D. rerio* embryos were imaged with a Stereo Lumar.V12 fluorescence microscope (Zeiss) equipped with GFP (Excitation 470/40nm; Emission 525/50nm and mCherry (Excitation 550/25nm; Emission 605/70nm) filters.

### Intestinal organoid culture

#### Establishment and maintenance

Crypts were isolated from the proximal part of the small intestine as reported previously^16^ and embedded in 15 μl droplets of BME-R1 (R&D Systems, 3433010R1) in a 48-well plate (Sigma, CLS3548-100EA). Organoids were established in WENR+Nic medium consisting of advanced DMEM/F12 (Gibco, 12634028) supplemented with penicillin/streptomycin (100x; Sigma, P0781), 10 mM HEPES (Gibco, 15630056), Glutamax (100x; Gibco, 35050061), B27 (50x; Life Technologies, 17504044), Wnt3 conditioned medium (Wnt3a L-cells, 50% of final volume), 50 ng/ml recombinant mouse epidermal growth factor (EGF; Gibco, PMG8041), 100 ng/ml recombinant murine Noggin (PeproTech, 250-38), R-spondin-1 conditioned medium (HA-R-Spondin1-Fc 293T cells, 10% of final volume) and 10 mM nicotinamide (MilliporeSigma, N0636). For the first week of culture, 100 μg/ mlprimocin (InvivoGen, ant-pm-05) and 10 μM ROCK-inhibitor/ Y-27632 (Sigma, Y0503) were supplemented to prevent microbial contamination and apoptosis, respectively. After the first passage, established organoids were converted to ENR budding organoid culture. Organoids were passaged at a ratio of 1:6 using mechanical dissociation.

#### 4-Hydroxytamoxifen treatment

Budding organoids at passage 3 or higher were passaged by mechanical dissociation and seeded in BME droplets. Medium containing vehicle (ethanol) or 500 nM 4-hydroxytamoxifen (Sigma, H7904) was added after BME polymerized. 8 hours later, medium was exchanged back to ENR and replenished every two days.

### Histology and immunohistochemistry

Samples were processed using a standard tissue protocol on an Automatic Tissue Processor Donatello (Diapath). Samples were embedded in paraffin and cut into 2 μm sections onto glass slides. Hematoxylin and eosin staining was carried out according to the standard protocol using a Gemini AS stainer (Thermo Scientific). For immunohistochemistry, the following antibodies were used: rabbit anti-Ki67 (1:200; 2 hours at room temperature; Abcam, Ab16667), rabbit anti-β-catenin (1:300; 1 hour at room temperature; Abcam, ab32572). For signal detection, a two-step HRP conjugated rabbit polymer system (DCS, PD000POL-K) was used. Stained slides were imaged with a 40x objective using the Pannoramic FLASH 250 III scanner (3DHISTECH) and images were cropped using the CaseViewer software.

### Immunofluorescence of intestinal organoids

BME droplets containing organoids were carefully collected into 1.5 ml tubes and spun down in a tabletop centrifuge at 600g for 5 minutes. The supernatant and visible fraction of attached BME were removed. The pellet was resuspended in 4% paraformaldehyde and fixed at room temperature for 15-20 minutes. Fixed organoids were washed 3 times in 1xPBS with 10-15-minute intervals. The organoids were blocked and permeabilized in a solution containing 5% DMSO, 0.5% Triton-X-100 (Sigma, T8787) and 2% normal donkey serum (Sigma, D9663) for one hour at 4 °C. The samples were stained overnight with Alexa 647-conjugated mouse anti-β-catenin (1:200; Cell Signaling Technology, 4627S) and ATTO 488-conjugated phalloidin (1:300; Sigma, 49409-10NMOL). The samples were washed three times with 1xPBS. During the last wash the samples were incubated with 2 μg/ml DAPI before mounting onto coverslips in a solution containing 60% glycerol and 2.5 M fructose^17^, followed by imaging on a multiphoton SP8 confocal microscope (Leica).

### Computing SCON targetable sites

In order to construct the databases for SCON insert sites, we used genomic information, about sequence, exon, coding region, and gene type, derived from Ensembl Biomart, build 102 (http://www.ensembl.org/)^18^ for mouse (*M. musculus*), rat (*R. norvegicus*), macaque (*M. mulatta*), marmoset (*C. jacchus*), and medaka (*O. latipes*). In order to map canonical transcripts to genes, we derived ‘PRINCIPAL:1’ from APPRIS database for mouse and rat with Ensembl, build 102 (http://appris-tools.org)^19^ and we regard the longest transcript as canonical transcript for other species. The quality features including GC-content, self-complementarity, and mismatch scores for each the candidate sites are mapped by same approach as CHOPCHOP (https://chopchop.cbu.uib.no)^20^. Of note, mismatch scores were calculated using Bowtie (v1.2.2)^21^ with ‘-v 3 -a’ option with 23 bp target sequences. The codes will be made available from the corresponding authors upon request.

## Acknowledgements

We thank members of Koo, Elling and Urban labs for valuable discussions and critical comments, Dr. Rike Zietlow for reading and correcting the manuscript, VBC core facilities (especially the Histopathology facility, BioOptics and the animal caretakers), the lab of Dr. Elly Tanaka for providing the frog embryos. This work was supported by core funding from the Institute of Molecular Biotechnology (IMBA) of the Austrian Academy of Sciences; ERC starting grant, Troy Stem cells, 639050; Interpark Bio-Convergence Center Grant Program; and fellowships to S.W. and K.G. (DOC Fellowship of the Austrian Academy of Sciences) and G.C. (Lise Meitner Postdoctoral fellowship M 2976, FWF).

## Author contributions

S.W. and B.-K.K. planned and designed the experiments. R.B., N.H., A.B., Y.Z. and S.W. cloned the constructs. R.B., A.B. and S.W. performed transfection experiments in Hek293T cells. R.B. performed experiments mouse ES cells. S.W. performed transfection experiments in C6, LLC-MK2, PK-15 and Vero cells. G.C. performed frog embryo injections. K.G. and A.P. performed zebrafish embryo injection. S.W. and B.-K.K. designed constructs for generating SCON mice. C.T. performed mouse embryo injection and subsequent procedures to generate SCON mouse lines. S.W. and N.H. performed mouse organoid experiments. S.W., S.P.C. and B.-K.K. performed mouse experiments and subsequent analysis. H.L. performed computational analyses for SCONable sites across different species. J.K., N.H., Y.Z. and S.P.C. performed supportive experiments. S.W. and B.-K.K. wrote the manuscript with input from the other authors.

## Declaration of Interests

The authors declare no competing interests.

## Supplementary information

**Supplementary Table 1**. List of CRISPR/Cas9-targetable exons for SCON insertion via MAGR sequence in *Mus musculus* genome.

**Supplementary Table 2**. List of CRISPR/Cas9-targetable exons for SCON insertion via MAGR sequence in *Rattus norvegicus* genome.

**Supplementary Table 3**. List of CRISPR/Cas9-targetable exons for SCON insertion via MAGR sequence in *Macaca mulatta* genome.

**Supplementary Table 4**. List of CRISPR/Cas9-targetable exons for SCON insertion via MAGR sequence in *Callithrix jacchus* genome.

**Supplementary Table 5**. List of CRISPR/Cas9-targetable exons for SCON insertion via MAGR sequence in *Oryzias latipes* genome.

**Supplementary Table 6**. List of CRISPR/Cas9-targetable exons for SCON insertion via VDGN sequence in *Mus musculus* genome.

**Supplementary Table 7**. List of CRISPR/Cas9-targetable exons for SCON insertion via VDGN sequence in *Rattus norvegicus* genome.

**Supplementary Table 8**. List of CRISPR/Cas9-targetable exons for SCON insertion via VDGN sequence in *Macaca mulatta* genome.

**Supplementary Table 9**. List of CRISPR/Cas9-targetable exons for SCON insertion via VDGN sequence in *Callithrix jacchus* genome.

**Supplementary Table 10**. List of CRISPR/Cas9-targetable exons for SCON insertion via VDGN sequence in *Oryzias latipes* genome.

**Extended Data Fig. 1.**
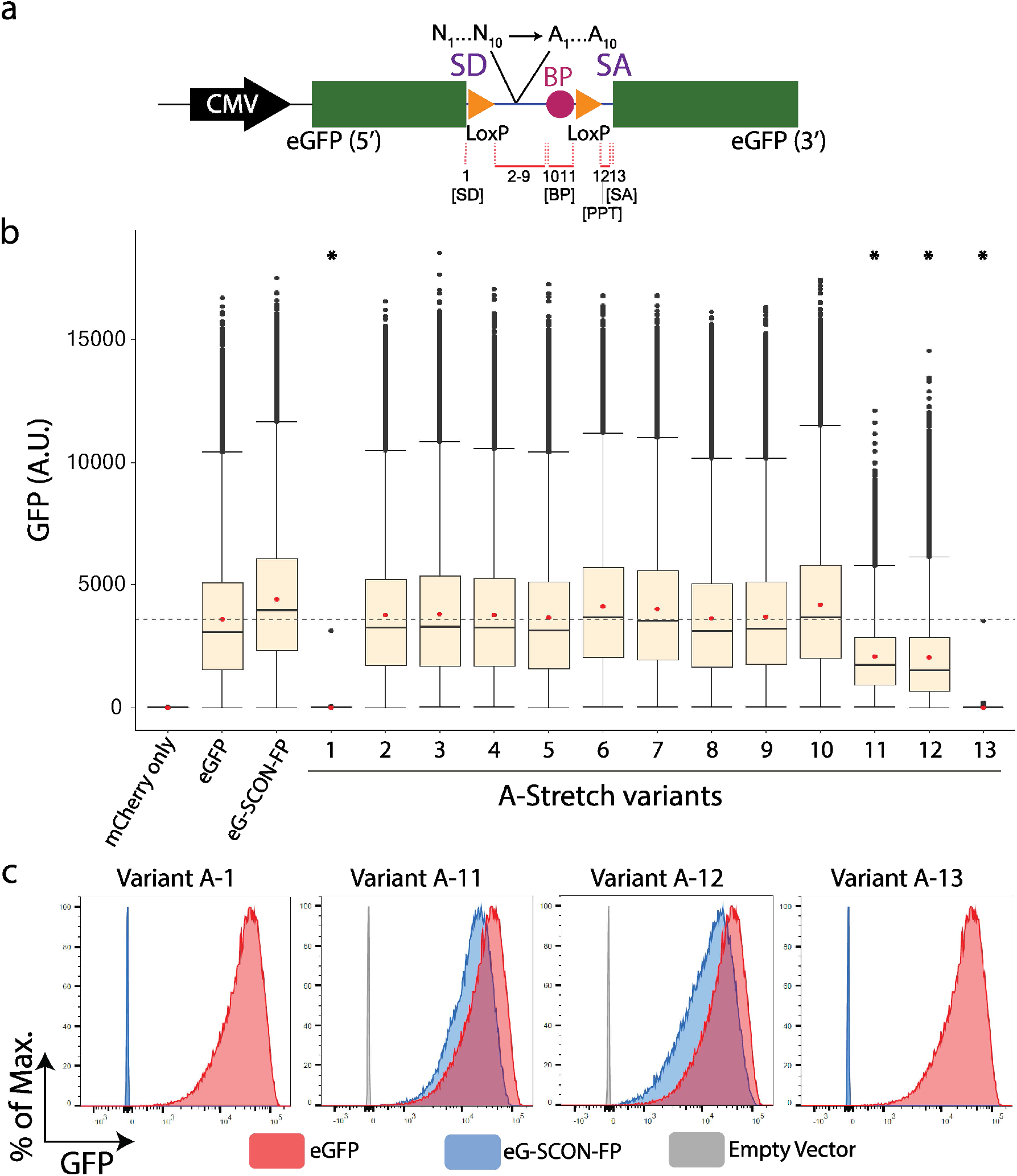
Dissection of functional sequences within the SCON cassette through A-stretch mutagenesis. **a**, Schematic diagram of the 13 A-stretch variants, where 6-10 nucleotides are converted to adenine, which cover all sequences from SD to SA, excluding the LoxP sites. PPT, polypyrimidine tract. **b**, Boxplots of measured scaled values from flow cytometry analysis of HEK293T cells transfected with mCherry only, eGFP, eG-SCON-FP and the 13 A-stretch variants. The red dot indicates the mean. The grey horizontal dotted line indicates the mean value of eGFP. **\*** indicates statistical significance (p = 0) from unpaired t-test with negative mean difference values when compared with the mean intensity value of eGFP. **c**, Flow cytometry of HEK293T cells transfected with either variant A-1, A-11, A-12 or A-13 (blue) compared with intact eGFP (red) and empty vector or mCherry only (grey).

**Extended Data Fig. 2.**
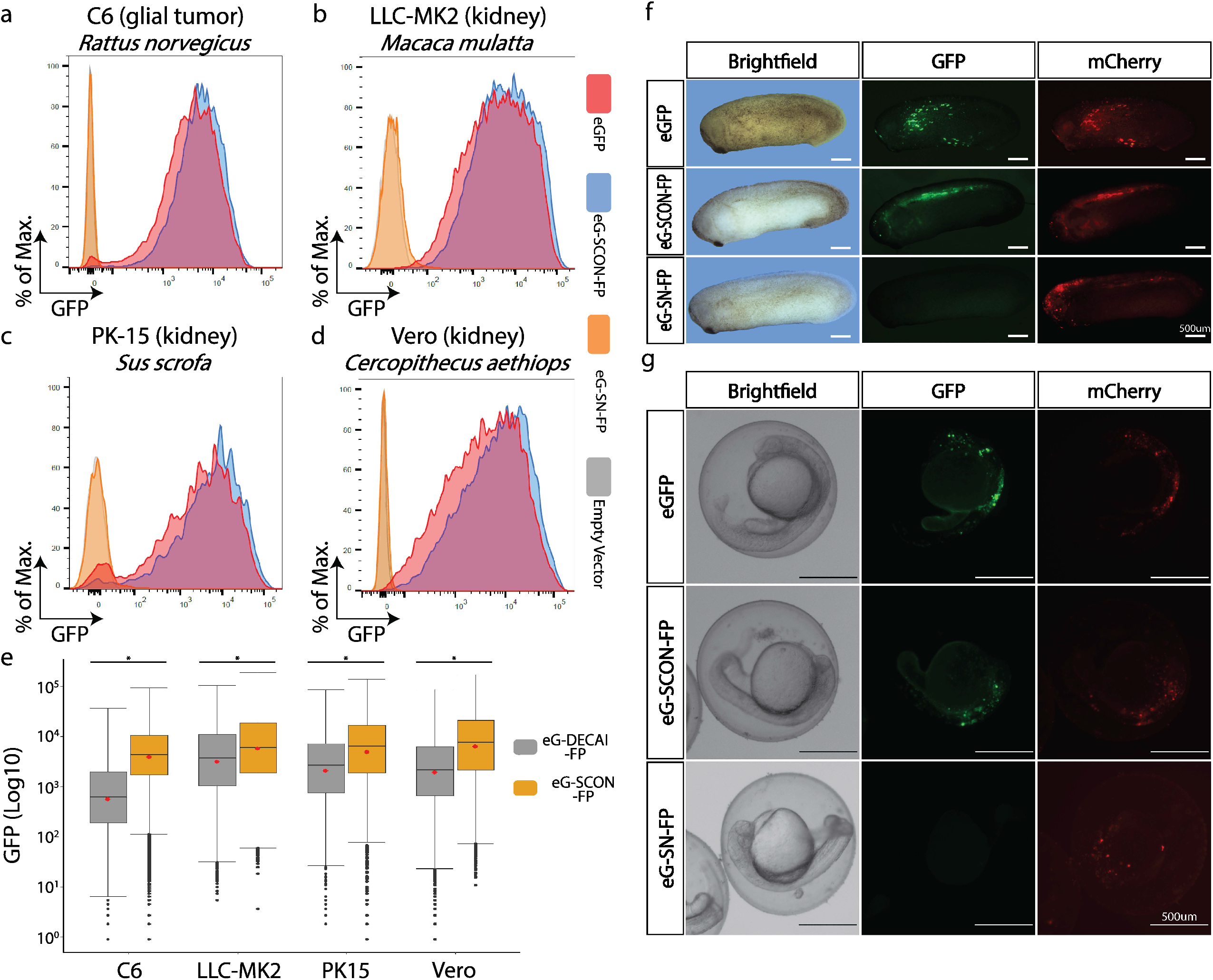
SCON is applicable and neutral in various vertebrate species. **a**, Flow cytometry analysis comparing the GFP levels in various cell lines transfected with eG-DECAI-FP (Grey) or eG-SCON-FP (yellow). *p<0.001, from unpaired t-test. **b-e**, Flow cytometry analysis of C6 (**b**), LLC-MK2 (**c**), PK-15 (**d**) and Vero (**e**) cells transfected with either eGFP (red), eG-SCON-FP (blue), eG-SN-FP (yellow) or empty vector/ mCherry only (grey). **f, g**, *Xenopus* and zebrafish embryos injected with eGFP, eG-SCON-FP or eG-SN-FP constructs, which were imaged 24 hours post-injection, a plasmid containing mCherry was used as an injection control. Scale bar, 500 μm.

**Extended Data Fig. 3.**
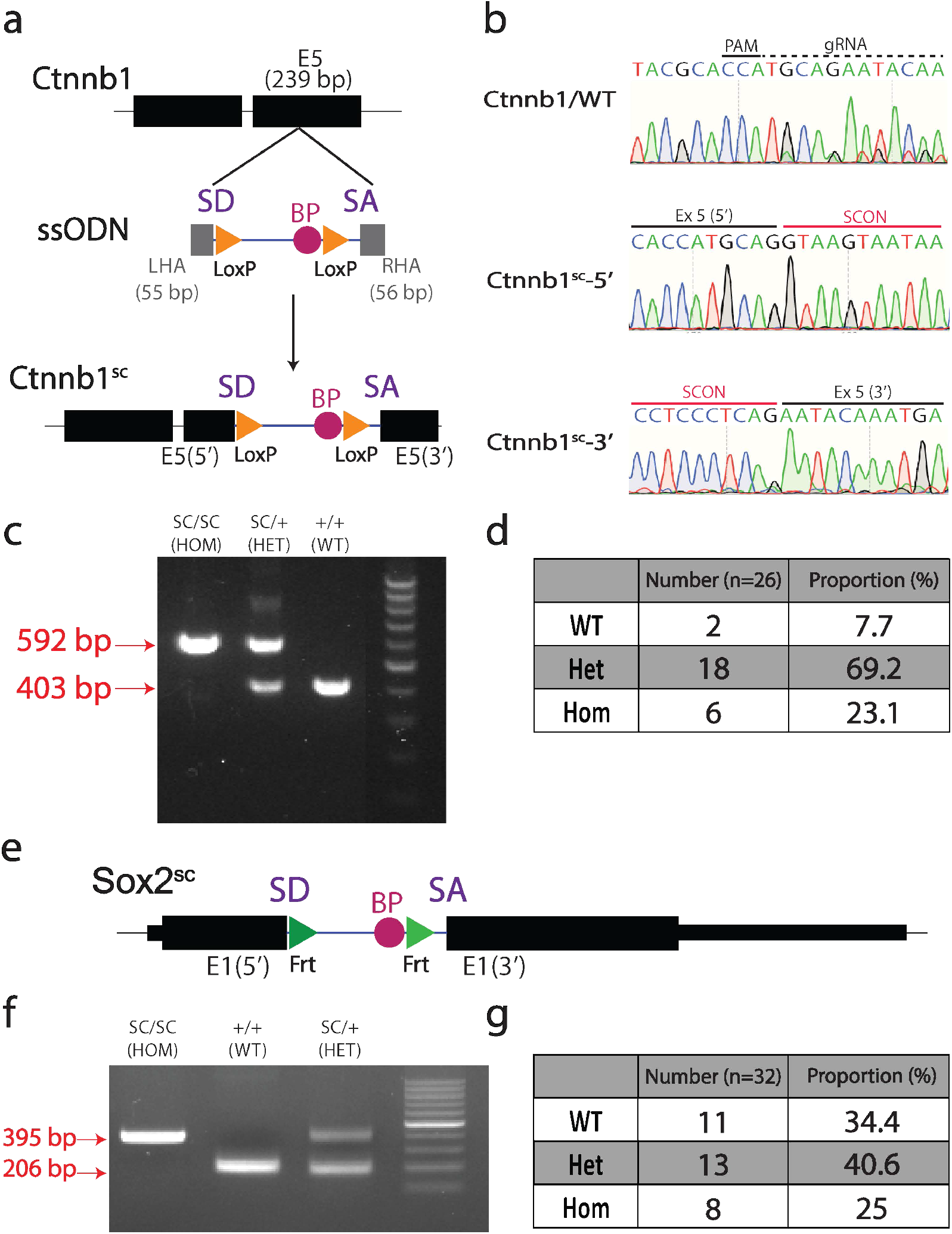
Ctnnb1^sc^ mouse generated via one-step embryo injection and is tolerated at homozygosity. **a**, Schematic drawing of SCON targeting ssODN, with 55 and 56 bp left and right homology arms, respectively, into the exon 5 of Ctnnb1 gene. **b**, Sanger sequencing track of the WT, 5’ and 3’ alleles of *Ctnnb1* and *Ctnnb1*^*sc*^, respectively. **c**, Genotyping PCR of *Ctnnb1*^*sc/sc*^ (HOM), *Ctnnb1*^*+/sc*^ (HET) and *Ctnnb1*^*+/+*^ (WT), of which the lower (403 bp) and upper (592 bp) bands correspond to the WT and knock-in alleles, respectively. **d**, Genotype quantification from crossings of double heterozygotes (*Ctnnb1*^*+/sc*^). Total number of offspring, n = 26. **e**, Schematic drawing of the *Sox2*^*sc*^ allele. **f**, Genotyping PCR of the Sox2^sc/sc^ (HOM), Sox2^+/+^ (WT) and Sox2^+/sc^ (HET), of which the lower (206 bp) and upper (395 bp) bands correspond to the WT and knock-in alleles, respectively. **g**, Genotyping quantification from crossings of heterozygotes (*Sox2*^*+/sc*^). Total number of offspring, n=32.

**Extended Data Fig. 4.**
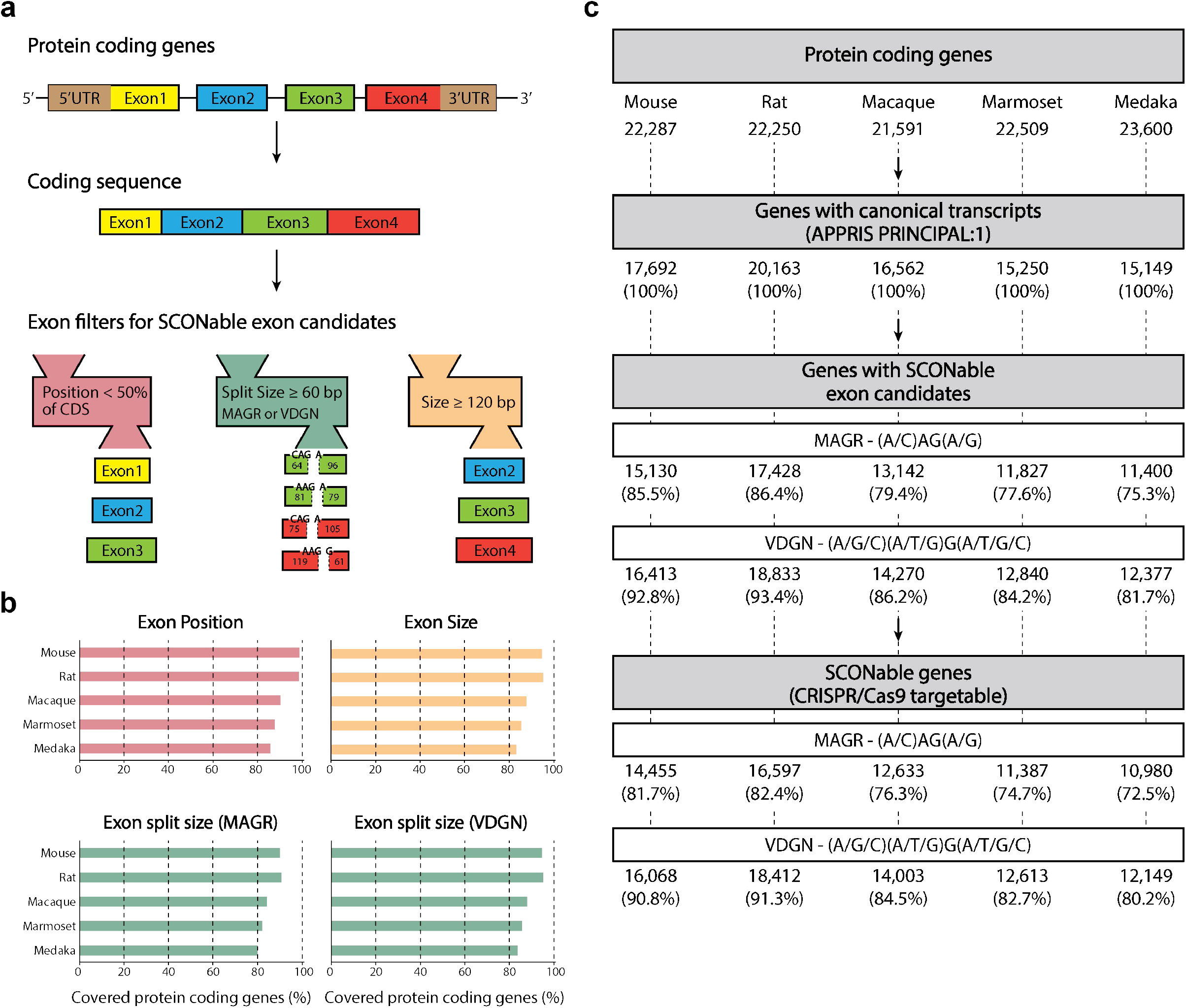
Scheme for construction of database for SCON insertion sites. **a**, Selection of the target exon candidates from protein coding transcripts. **b**, Gene coverages of individual exon filters such as position, size, and split exon size. **c**, Summary of databases for SCON insertion sites with CRISPR/Cas9 targeting sites.

## Notes

### Competing Interest Statement

The authors have declared no competing interest.

